# Photolipid excitation triggers depolarizing optocapacitive currents and action potentials

**DOI:** 10.1101/2023.08.11.552849

**Authors:** Carlos A. Z. Bassetto, Juergen Pfeffermann, Rohit Yadav, Simon Strassgschwandtner, Toma Glasnov, Francisco Bezanilla, Peter Pohl

## Abstract

Optically-induced changes in membrane capacitance may regulate neuronal activity without requiring genetic modifications. Previously, they mainly relied on sudden temperature jumps due to light absorption by membrane-associated nanomaterials or water. Yet, nanomaterial targeting or the required high infrared light intensities obstruct broad applicability. Now, we propose a very versatile approach: photolipids (azobenzene-containing diacylglycerols) mediate light-triggered cellular de- or hyperpolarization. As planar bilayer experiments show, the respective currents emerge from millisecond-timescale changes in bilayer capacitance. UV light changes photolipid conformation, which awards embedding plasma membranes with increased capacitance and evokes depolarizing currents. They open voltage-gated sodium channels in cells, generating action potentials. Blue light reduces the area per photolipid, decreasing membrane capacitance and eliciting hyperpolarization. If present, mechanosensitive channels respond to the increased mechanical membrane tension, generating large depolarizing currents that elicit action potentials. Membrane self-insertion of administered photolipids and focused illumination allows cell excitation with high spatiotemporal control.

**Highlights:**

- Rapid photolipid photoisomerization generates optocapacitive currents in planar lipid bilayers and HEK293 cells.
- These currents originate from photo-induced changes in membrane capacitance
- UV light-triggered membrane depolarization opens Na_V_1.3, evoking action potentials.
- Blue light-induced mechanosensitive channel opening gives rise to depolarizing currents, which may evoke Na_V_1.3-mediated action potentials.

---

Regulating neuronal activity by light has sparked scientists’ interest for several decades ^1^. By allowing for tight spatiotemporal regulation, light-based approaches – notably optogenetics – aid in deciphering neuronal networks, supplementing traditional electrical techniques ^2, 3^. Optogenetic approaches frequently rely on light-gated cation-conducting channels, e.g., channelrhodopsins ^4^. Exposing a channelrhodopsin-harboring neuron to light increases membrane cation permeability which causes depolarization and, eventually, neuronal spiking ^5^. Effectively, this technique allows for millisecond-timescale and cell-type-specific optical control of neurons ^6^. Yet, given that mammalian neurons do not generally express light-gated cation channels, genetic material encoding these proteins has to be transfected and subsequently expressed by the target cells ^7^.

Photothermal optocapacitive approaches circumvent the necessity for introducing exogenous genetic material ^8-10^. These techniques are based on a rapid light-evoked increase in cellular membrane temperature, leading to a fast rise in bilayer capacitance that causes a depolarizing optocapacitive current. Mechanistically, heating lipid bilayers causes them to expand in area and thin, entailing the required increase in capacitance ^11, 12^. The rate of change of capacitance determines action potential (AP)-generating effectiveness and critically depends on the rate of temperature change ^10, 13^. Membrane-associated extrinsic nanomaterials that convert visible light into heat ^9^, and high-energy pulses of infrared light absorbed by water in the membrane vicinity ^8^ have achieved a sufficiently fast increase in membrane temperature for photothermal optocapacitive depolarization and consequent AP generation.

Photosensitive molecules represent yet another approach for regulating neuronal activity by light ^14^. They can be classified by the mechanism of interaction with their target – non-covalent or covalent binding – as well as by the reversibility of their action ^14^. For example, photoswitchable molecules covalently bound to glutamate receptors served to evoke or terminate trains of action potentials ^15^. Non-covalently bound light-sensitive diacylglycerols with two azobenzene-containing acyl chains (OptoDArG) were used to regulate cation permeation and membrane potential by opening TRPC3 channels ^16^. Albeit such photolipids may act according to a lock-and-key principle, as in the aforementioned examples, by partitioning into biological membranes and changing their molecular structure upon photoisomerization, they affect lipid bilayer material properties such as thickness, surface area, bending rigidity, compressibility and the propensity towards lipid domain formation ^17-22^. It is well established that these properties may contribute to the gating of mechanosensitive and mechanically-modulated ion channels ^23-26^. Photoisomerization-induced changes in bilayer material properties and tension may alter channel open probability or dynamics of membrane proteins ^19, 27, 28^.

Recently, neuronal excitation was achieved using the membrane-partitioning azobenzene-containing photoswitchable compound Ziapin2 ^29^. The photoisomerization of trans-Ziapin2 by blue light led to a rapid decrease in membrane capacitance. Conceivably, this drop produced a hyperpolarizing capacitive current. Yet, conductances of other origins also contributed as the observed hyperpolarizing current (i) did not linearly depend on voltage and (ii) persisted nearly unaltered for ≈250 ms even though the capacitance change took no longer than ≈20 ms ^30^. Subsequently, delayed depolarization ensued by an elusive mechanism. Being unaware of the channels that might have facilitated the depolarizing current, the authors speculated that Ziapin2 caused a spontaneous fast capacitance increase (*t*_1/2_≈0.2 s). However, Ziapin2’s unprompted return to its cis conformation in DMSO takes orders of magnitude longer (*t*_1/2_=108 s) ^30^.

Here, we used minimal systems to identify the molecular mechanisms underlying photolipid-induced hyperpolarization and depolarization. We first used planar lipid bilayers to observe UV light-triggered depolarizing currents and subsequently exploited them to trigger APs by photolipid photoisomerization in sodium channel (Na_V_1.3)-expressing HEK cells. We also clarified that mechanosensitive channels present in HEK cells may facilitate the depolarizing currents required for AP generation upon blue light-triggered hyperpolarization. We base this study on the photolipid OptoDArG (Fig. 1a), as diacylglycerols are (i) ubiquitous in biological systems, (ii) known to alter the mechanical properties of lipid bilayers upon photoswitching if equipped with azobenzene moieties ^19^, and (iii) able to spontaneously flip-flop across membranes which ensures their presence also in the inner leaflet of plasma membranes when added from the outside medium.

**Fig. 1:**
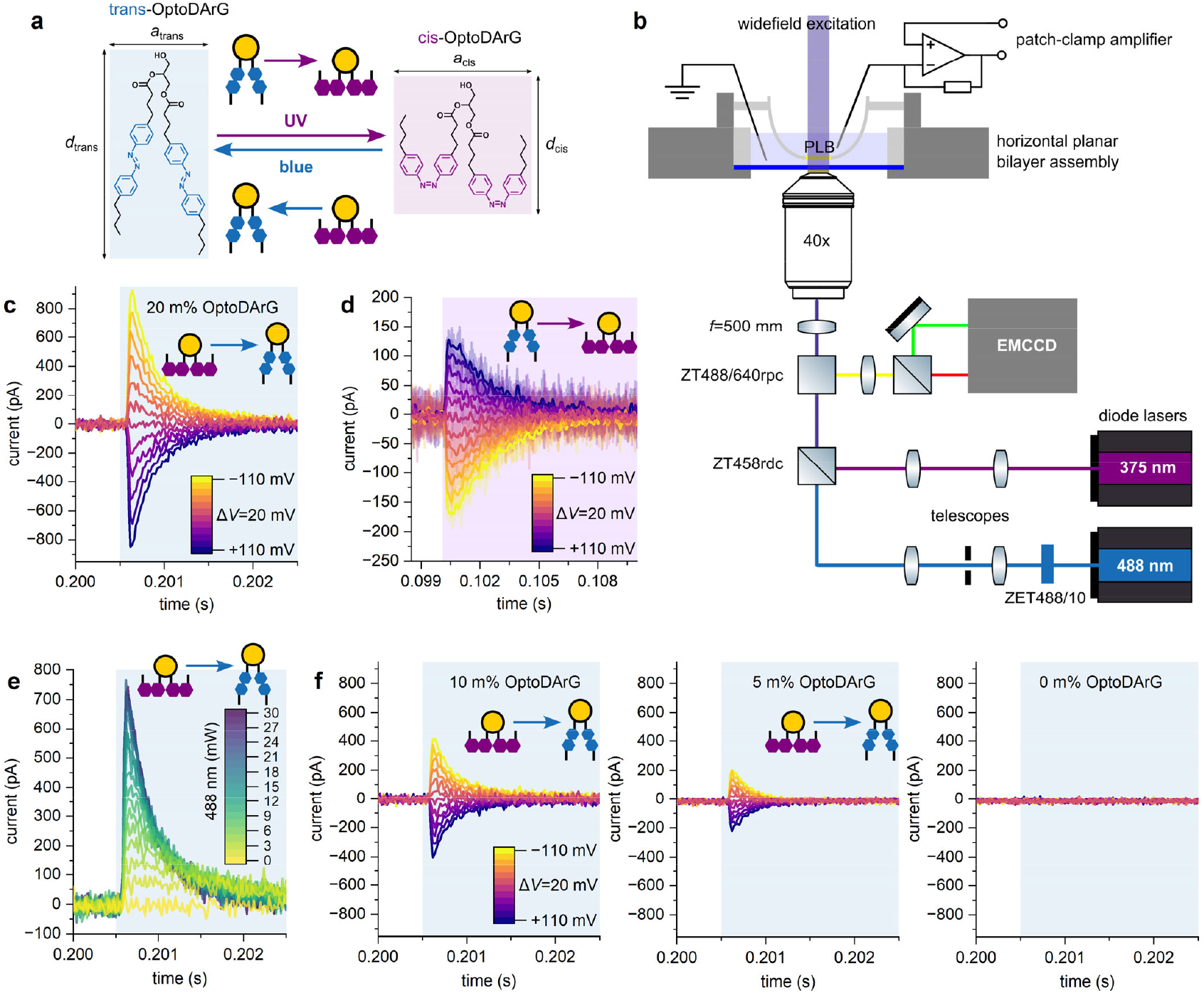
Generation of hyper- and depolarizing optocapacitive currents in PLBs. **a**, Rationale for capacitance changes upon OptoDArG photoisomerization: cis-OptoDArG is a broader (*a*_cis_>*a*_trans_) and shorter (*d*_cis_<*d*_trans_) molecule than trans-OptoDArG. Upon incorporation into bilayers, these differences in molecular structure alter bilayer geometry upon photoisomerization: UV light favors cis-OptoDArG, which increases bilayer surface area and reduces thickness. Blue light achieves opposite effects. According to Eq. 1, these geometrical alterations translate into differences in capacitance. **b**, A schematic representation of the horizontal PLB setup (see Methods). **c**, Optocapacitive currents were generated upon rapid photoisomerization of membrane-embedded cis-OptoDArG by blue light (blue background). The PLB was clamped at voltages ranging from −110 mV to +110 mV with Δ*V*=20 mV, as indicated by the color legend. Current traces obtained at different *V* are overlaid. The laser was kept on for 50 ms to ensure quantitative switching whereby the evoked currents decayed to electrical noise level within milliseconds. **d**, Recorded as **c**, but the PLB was in its trans photostationary state prior to UV light exposure (purple background). The raw currents are overlaid with smoothed curves for clarity. **e**, Overlay of optocapacitive currents recorded at *V*=−110 mV and different blue light intensity (the numbers in the color legend give power at the sample stage in mW). **f**, To demonstrate the dependence of *I*_cap_ on the fraction of OptoDArG in the PLB-forming lipid mixture (see Eq. 7), the latter was reduced from 20 m% (as in **c** to **e**) to 10 m% and 5 m%. At 0 m%, blue light exposure evokes no currents. The recordings were done as in **c**.

## Photolipid excitation elicits optocapacitive currents

Differences in molecular structure between OptoDArG’s photoisomers affect the geometry of bilayers containing them: the extended cis-OptoDArG conformation awards bilayers with increased surface area, *A*, and reduced thickness of the membrane’s hydrophobic core, *d*_hc_, whilst its trans isomer requires less surface area per molecule and increases *d*_hc_ (Fig. 1a) ^19^. Consequently, photoisomerization of membrane-embedded OptoDArG changes lipid bilayer capacitance, *C*:

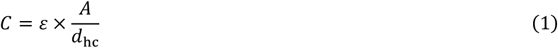

where *ε* denotes absolute permittivity. For achieving optocapacitive modulation of the membrane potential, switching *C* within tens of seconds to minutes as in our earlier study ^19^ would not suffice. As reflected by Eq. 2 the rate of change of *C*, d*C*/d*t*, determines the AP-generating effectiveness by the optocapacitive approach ^10, 13^, i.e., the amplitude of the optocapacitive current, *I*_cap_:

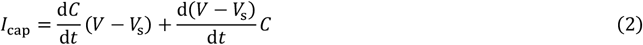

where *V* and *V*_s_ denote transmembrane and membrane surface potential differences ^10, 11, 13^. Under voltage-clamp conditions d*V*/d*t*=0, and Eq. 2 simplifies to:

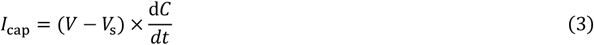

Eq. 3 assumes that *V*_s_ remains unaltered during photoswitching. The condition is undoubtedly fulfilled in a planar lipid bilayer (PLB) with symmetrical leaflets, since *V*_s_≈0 mV. For the plasma membrane of *E. coli* with *V*_s_=−18 mV ^31^, a 5% change in the area per lipid would alter *V*_s_ by no more than 1 mV, which is negligible.

To evoke *I*_cap_ by photolipid photoisomerization, we folded horizontal solvent-depleted PLBs (diameter typically 60– 80 μm) from 80 m% *E. coli* polar lipid extract (Avanti Polar Lipids) and 20 m% OptoDArG and placed them within working distance of the 40× magnification objective of an inverted widefield fluorescence microscope (Fig. 1b). Eq. 3 predicts *I*_cap_ in the range of tens to hundreds of picoampere for *V*=100 mV and a 2.5 pF change in *C* that occurs within a few milliseconds. The required time depends on the rate of isomerization (see below).

Before exposure to blue laser light (≈30 mW, ≈58 μm 1/e²-diameter, 488 nm), a UV laser diode (375 nm) switched OptoDArG into a cis-enriched photostationary state. Blue light illumination triggered positive *I*_cap_ when we clamped the PLB at negative *V*, and negative *I*_cap_ for positive *V* (Fig. 1c). Since photoisomerization from cis-to trans-OptoDArG by blue light reduces *C*, this result is in line with Eq. 3. Further, this observation rules out a photothermal mechanism induced by blue light because heating mandates d*C*/d*t*>0 which would generate oppositely-directed *I*_cap_ ^8, 10, 11^. Subsequent rapid switching of the PLB into a cis-enriched photostationary state using UV laser light (≈30 mW incident at a diameter of roughly 150–250 μm) generated positive *I*_cap_ at positive *V* (Fig. 1d). Again, this is consistent with Eq. 3 since cis-OptoDArG increases *C* and, thus, d*C*/d*t* is positive.

To determine the factors governing *I*_cap_, we begin by expressing the time course of *C* upon blue, *C*_blue_, and upon UV light exposure, *C*_UV_, as follows:

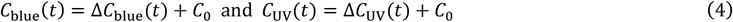

where *C*_0_ is *C* at the onset of light exposure, Δ*C*_blue_ the decrement in *C* upon blue light exposure, and Δ*C*_UV_ the increment in *C* upon UV light exposure. Changes in *C* upon photoisomerization emerge from the structural transition of many individual photolipids. As part of the bilayer capacitor, each cis and trans photolipid contributes to *C*, and we refer to the individual contributions as *c*_c_ and *c*_t_; conceivably, *c*_c_>*c*_t_. Consequently, Δ*C*_blue_ and Δ*C*_UV_ depend on the number of cis- and trans-OptoDArG molecules in the bilayer *n*_c_ and *n*_t_:

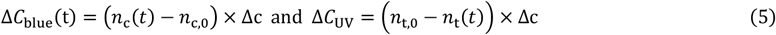

whereby Δ*c*=*c*_c_−*c*_t_. *n*_c,0_ and *n*_t,0_ denote the number of cis and trans photolipids at time *t=*0 s. *n*_c_ and *n*_t_ decay depending on blue, *I*_blue_, and UV light irradiance, *I*_UV_, as:

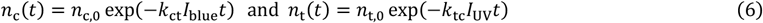

where *k*_ct_ and *k*_tc_ (in m^2^W^−1^s^−1^) denote rate constants of OptoDArG photoisomerization from cis to trans and trans to cis. Though of the same order of magnitude, *k*_ct_ and *k*_tc_ adopt different values ^32^. They may be considered to be independent of power ^32^. Also, Eq. 6 considers *k*_tc_ upon blue and *k*_ct_ upon UV light exposure to be zero, which, albeit a good approximation, is quantitatively incorrect due to overlaps in photoisomer absorption ^32, 33^. Eq. 6 also assumes a negligibly small contribution of thermal relaxation of the metastable cis photoisomer at the millisecond timescale ^22^. From Eqs. 3 and 6 we find:

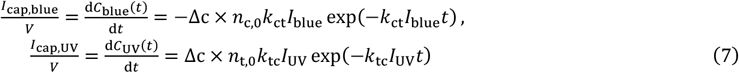

where *I*_cap,blue_ and *I*_cap,UV_ denote optocapacitive currents evoked upon blue and UV light exposure.

For small sizes of the solvent torus anchoring the PLB and millisecond intervals, we may assume that *n*_c_+*n*_t_ is constant, i.e., lipid trafficking between the bilayer and the PLB’s lipid reservoirs (torus and monolayers) is negligible. Then Eq. 7 predicts an exponential decay of *I*_cap_ – which we tentatively observe in Figs. 1c and 1d. Indeed, we find that the power-dependent *I*_cap,blue_ in Fig. 1e are well-grasped by simple exponential decay fits (Supplementary Fig. 1a). The obtained monoexponential rate constants *k*_blue_=*k*_ct_*I*_blue_ are linear with irradiance up to ≈ 750 Wcm^-2^ blue light with 1/*k*_blue_ in the millisecond range. *k*_blue_ levels off abruptly at >750 Wcm^-2^, conceivably due to instrumental (filtering) and fitting limitations (Supplementary Fig. 1b). 1/*k*_blue_ is orders of magnitude larger than the time (ps) required for switching individual azobenzene moieties ^34^ because the absorption probability is limiting. A linear model fit reveals a slope of *k*_ct_ = 3.85×10^−3^ cm²mW^−1^s^−1^ (R²>0.99). We note that our blue illumination profile was not flat-top but Gaussian, so we estimate irradiance by calculating the area from the 1/e²-diameter of the illumination profile (≈58 μm). *k*_ct_ is reasonably close to the rate of photoisomerization of azobenzene-containing surfactants of 3−4×10^−3^ cm²mW^−1^s^−1^ at 490 nm ^32^. This agreement indicates that the observed *I*_cap_ values are an immediate consequence of changes in *C* due to photolipid photoisomerization. As also predicted by the model, *k*_blue_ and *k*_UV_ are independent of *V* (Supplementary Fig. 2).

In contrast to the prediction of Eq. 7 and the experimental results in PLBs shown in Supplementary Fig. 3a, a previous report claiming to have observed optocapacitive currents ^29^ displayed light-triggered currents that persisted nearly unaltered as long as light exposure lasted ^30^. The molecular origin of these persisting currents remained elusive. Perhaps the dimerization propensity of Ziapin2, the molecule with azobenzene moieties replacing photolipids in the latter study, is responsible for it. Alternatively, the activation of membrane ion channels may provide an explanation.

It is important to note that in absence of OptoDArG no optocapacitive currents is observed, further demonstrating that the effects observed in PLB containing OptoDArG are not light-induced thermal effects (Supplementary Fig. 3b).

Eq. 7 proposes two straightforward strategies to enhance *I*_cap,blue,_ and *I*_cap,UV_:

a. Increasing *I*_blue_ or *I*_UV_ ought to increase d*C*/d*t*. For demonstration, we clamped a PLB at *V*=−110 mV and repeatedly exposed it to blue light of increasing intensity (Fig. 1e).
b. As demonstrated in Figs. 1c and 1f, decreasing OptoDArG concentration, i.e., *n*_c,0_ and *n*_t,0_ in the PLB-forming lipid mixture from 20 m% to 5 m% reduces initial optocapacitive current amplitude.

### Light-induced capacitance changes generate *I*_cap_

Consecutive 50 ms blue (blue background) and UV (purple background) light exposures induced fast changes in *C* of PLBs (Fig. 2a, b). Zooming into the first milliseconds of the exposure times (Figs. 2c and 2d) reveals exponentially decaying capacitance traces. Differentiating the latter (Eq. 3) numerically by calculating difference quotients and multiplying the obtained derivative traces by *V*=±110 mV yielded current traces. Overlaying these calculated traces with *I*_cap_ (obtained under voltage-clamp at *V*=±110 mV) showed a reasonable match with the recorded current (Figs. 2e and 2f). The small residual deviations may be due to (i) the time delay between *I*_cap_ and *C* recordings and (ii) limitations set by the rate with which we can measure membrane capacitance. The electrodes and the deployed software lock-in amplifier (HEKA) limited us to 5 kHz sine frequency of the probing signal. Ideally, we would have aimed for a rate closer to 20–50 kHz.

**Fig. 2:**
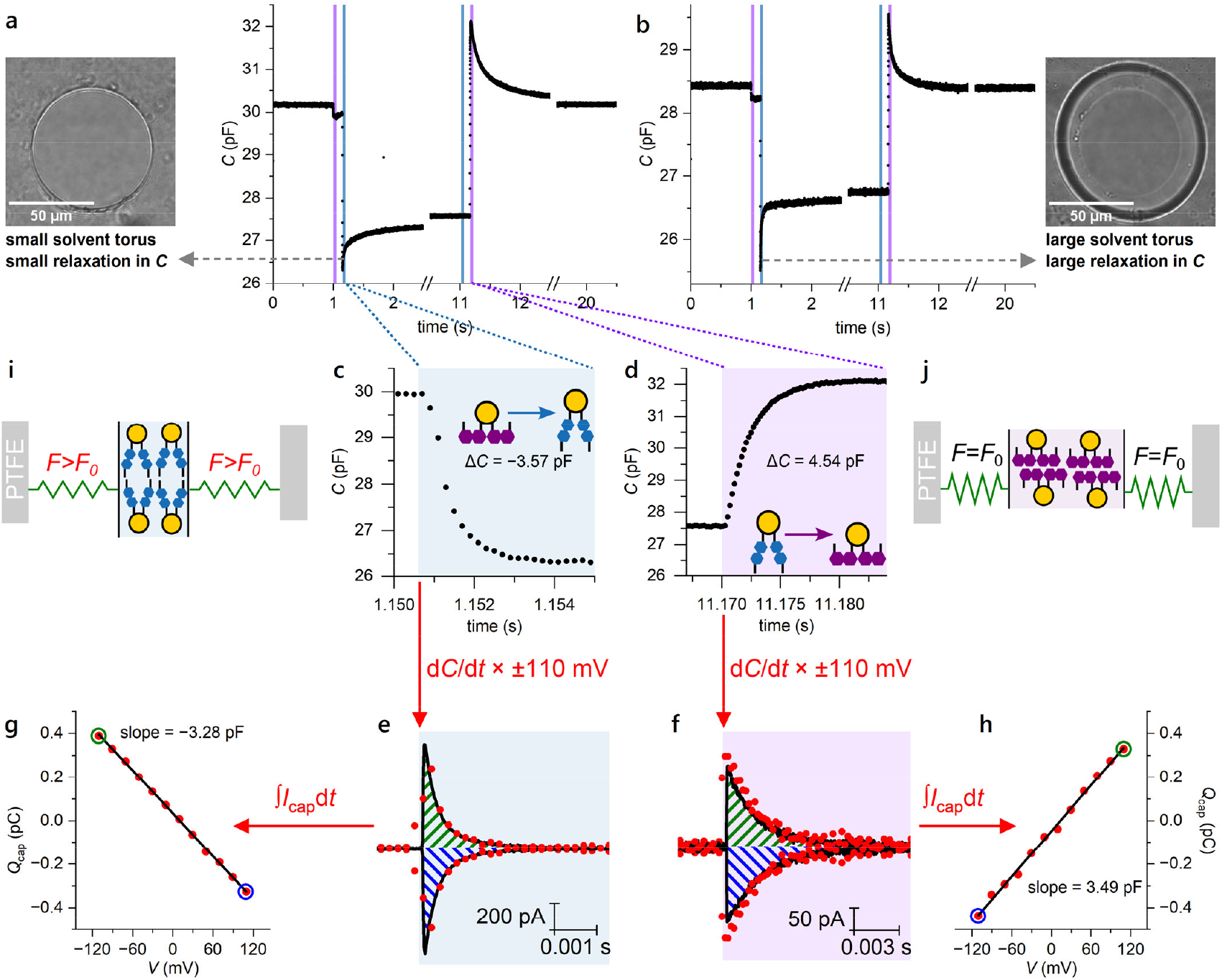
Light-induced capacitance changes generate *I*_cap_. **a**,**b**, Time trace of PLB capacitance (black line; an average of 5 consecutively-recorded traces) indicating the 50 ms exposure intervals to blue light (blue background) and UV light (purple background). UV illumination immediately prior to blue light exposure at 1.15 s and blue light illumination immediately prior to UV light exposure at 11.17 s ensures a cis- and trans-enriched photostationary state, respectively. In **a**, the torus of the solvent-depleted PLB was small (image on the left). In **b**, *C* was recorded in the presence of a large solvent torus, as apparent in the corresponding PLB image on the right. Note the increased relaxation amplitude following blue light exposure relative to **a. c, d**, *C* changes exponentially as zoom-ins into the (**c**) blue (blue background) and (**d**) UV light (purple background) illumination intervals shown in **a. e, f**, The graphs below the recordings of *C* show *I*_cap_ with (**e**) blue and (**f**) UV light exposure recorded at *V=*±110 mV (black lines; each averaged from 20 consecutively-recorded current traces). The overlaid red points are calculated from the respective above capacitance trace by numerical differentiation and subsequent multiplication with ±110 mV, as indicated by the arrows. **g, h**, The graphs show *Q*_cap_ obtained upon integration of *I*_cap_ in Fig. 1c (**g**) and Fig. 1d (**h**) plotted over *V*; acc. Eq. 8, the slope of a linear fit (black line) corresponds to Δ*C* upon photoisomerization (R²>0.99). The green and blue circles correspond to the areas marked with the same color in **e** and **f. i, j**, Schematics of how photoisomerization from cis- to trans-OptoDArG generates tension within the membrane. The grey bars labeled PTFE symbolize the septum aperture within which the PLB is mounted, and green springs refer to the bilayer lipids other than OptoDArG. Upon blue light exposure, the area per photolipid decreases (*a*_trans_<*a*_cis_) which, as the PLB is anchored laterally, causes stretching of the remaining lipids in the bilayer. The loading of the springs symbolizes tension within the PLB.

Vice versa, we note that the photoisomerization-induced Δ*C* can be found by integrating *I*_cap_ (Eq. 3):

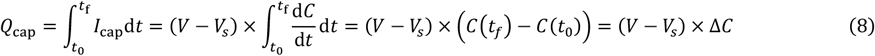

where *Q*_cap_ denotes the number of capacitive charges displaced upon photoisomerization, *t*_0_ the starting time of light exposure, and *t*_f_−*t*_0_ the integration time. To demonstrate this approach, we numerically integrated *I*_cap_ in Figs. 1c and 1d deploying an integration time of 3 ms and 10 ms, respectively. Figs. 2g and 2h show the resulting values for *Q*_cap_ plotted against *V*; according to Eq. 8, the slope of the indicated linear fits equals Δ*C*. We find that the thus obtained values for Δ*C* (−3.28 pF and +3.49 pF) underestimate those obtained from direct capacitance recordings (−3.57 pF (Fig. 2c) and +4.54 pF (Fig. 2d)) by ≈10–25%. These deviations partly originate from finite integration times and analog filtering of the current traces at 10 kHz.

An additional contribution to *I*_cap_ and Δ*C* comes from the membrane torus. We observe that in the presence of a small torus, the rapid blue light-mediated drop in *C* is followed by a fast relaxation in *C* (between 1.2 and 1.3 s in Fig. 2a, indicated by the gray arrow) that amounts to <15% of the amplitude of the drop. In all likelihood, the relaxation indicates a shape change of the torus due to a pulling force of the PLB. The force originates from area differences between cis and trans photolipids. If the torus is larger (Fig. 2b), this relaxation may reach >30%. We commonly discarded such traces. Interestingly, relaxation in *C* also follows the photolipid-mediated increase in *C* upon UV light exposure. Conceivably, it signifies that the constitutive tension of PLBs flattens photo-induced undulations that appear with the increment in membrane area upon membrane thinning. Importantly, these torus-specific relaxations do not represent an intrinsic feature of the photolipid. These minor drawbacks of the experimental model system notwithstanding, we find a general agreement between current and capacitance recordings. Applications of Eq. 3, Eq. 7 and Eq. 8 provide strong evidence for the optocapacitive mechanism of rapid photolipid photoisomerization.

### The transition to trans-OptoDArG generates membrane tension

Rapid photoisomerization from cis-to trans-OptoDArG generates membrane tension, as schematically depicted in Figs. 2i and j. First, exposure to blue light reduces area per photolipid (*a*_trans_<*a*_cis_). Second, the narrower photolipids exert a pulling force on the neighboring lipids and thus stretch them temporarily (illustrated by the green springs). Equivalently, one may consider that the reduction in photolipid area leads to a transient reduction in bilayer-internal lateral pressure due to less steric hindrance ^27^. The result is the same: tension components temporarily outweigh pressure components in the bilayer’s lateral pressure profile, and, as a result, membrane tension emerges ^35^.

As a back-of-the-envelope calculation shows, the achievable tension, τ, is relevant for mechanosensitive proteins. For example, mechanosensitive channels like MscL open at τ ≈10 mN/m ^36^. Such τ value translates for membranes with a typical stretching modulus, K, of 250 mN/m into an area increment of 4 % (α = 0.04), since τ = K α. The corresponding energy E per membrane area, A, required to stretch the membrane is equal to:

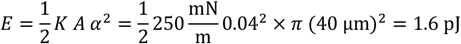

The work performed by isomerization of a single azobenzene switch has been estimated at 4.5×10^−20^ J ^37^. Assuming that a lipid molecule occupies, on average, 0.7 nm^2^, the number of photolipids (10 mass-% OptoDArG) per leaflet amounts to *π* (40 μm)^2^/(10 x 0.7 nm^2^) 7 10^8^, we find a maximal work of ≈ 63 pJ, the bilayer could perform. We conclude that photolipid switching may exert work for the opening of mechanosensitive channels, even if (i) less than 100 % of the photolipids change their conformation, (ii) the deformation of other lipids and flattening of membrane undulations contribute to the expenditure of *E*.

Eventually, the photo-induced tension will relax because additional lipids from the top of the septum and torus may be pulled into the PLB. The situation may be similar in cells as membrane invaginations may also provide a lipid reservoir, as previously observed with osmotically challenged cells ^38^. We demonstrate the photoactivation of mechanosensitive channels below to provide experimental evidence for photo-induced tension in the cellular membrane.

### Photolipid-evoked hyper- and depolarizing currents in cells

OptoDArG self-inserts into both leaflets of the plasma membrane when dissolved in DMSO and administered to the aqueous solution. For the cell experiments, we used a setup optimized for whole-cell patch-clamp experiments (Fig. 3a). The intensity of the Ti : Sapphire laser (367 nm after the second harmonic generator) was regulated by a polarizer. The pulse duration was controlled by a shutter that allowed the generation of millisecond pulses with a sharp rise time (less than 20 μs). An in-house designed and built uncoated fused silica objective optimized the intensity of UV light reaching the sample stage. As in PLBs, we found that exposing the cells (Fig. 3b) to millisecond UV and blue light pulses (445 nm) generated hyper- and depolarizing currents, respectively (Fig. 3c). In accordance with Eq. 7 and in line with our PLB recordings (e.g., Fig. 1e), the optocapacitive currents in Fig. 3c decay exponentially. Importantly, no optocapacitive current was observed when cells were not labeled with OptoDArG (Supplementary Fig. 4a), ruling out the possibility that the currents shown in Fig. 3c were thermally evoked.

**Fig. 3:**
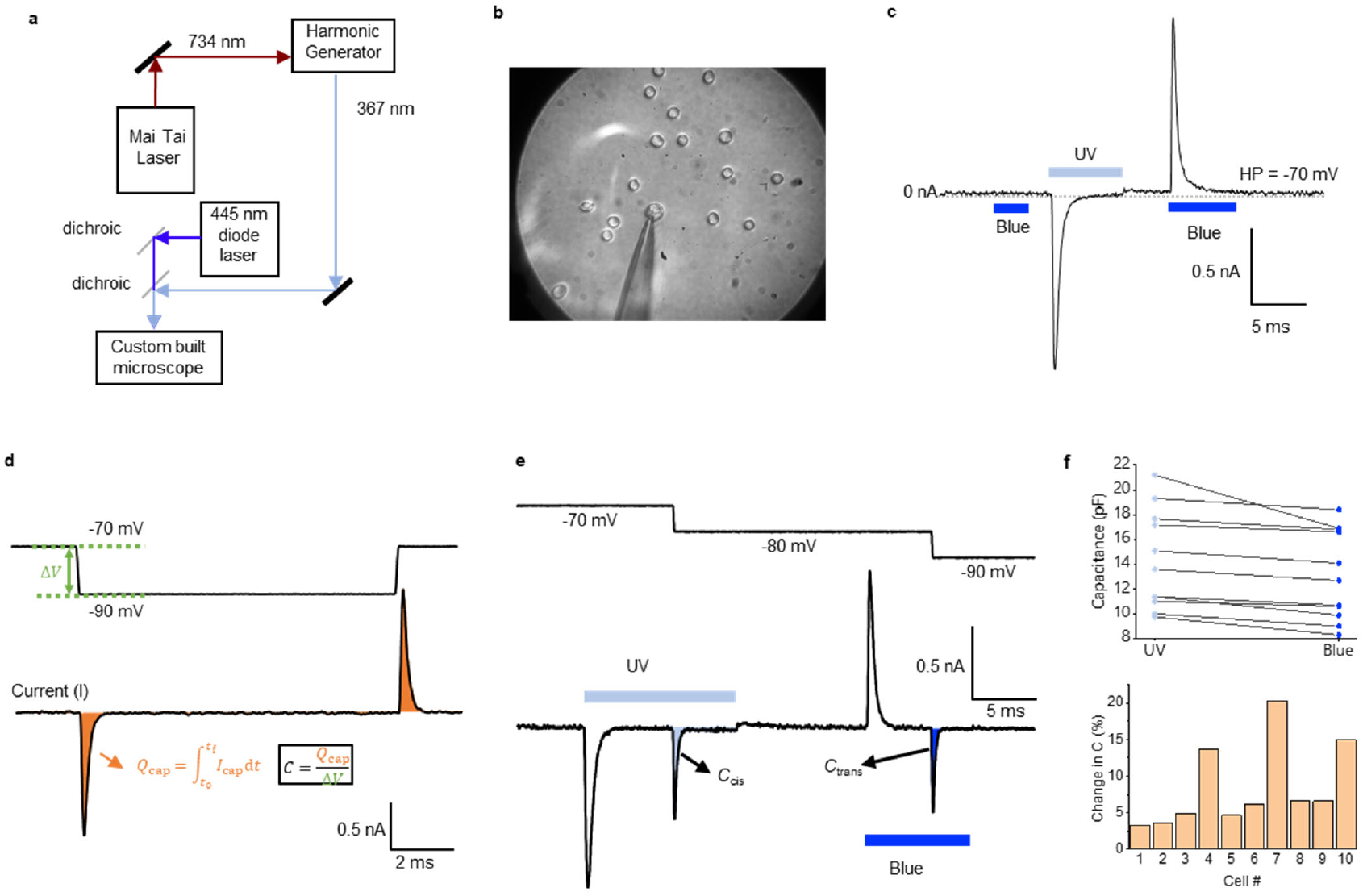
OptoDArG photoisomerization generates hyper- and depolarizing capacitive currents in HEK293 expressing Na_V_1.3. **a**, Schematic representation of the cell stimulation setup. **b**, HEK cells were labeled with 60 μM OptoDArG for 1 hour (the labeling protocol is described in the Methods section). The image was taken through the in-house designed fused silica objective. **c**, Whole-cell voltage-clamp recordings on HEK293 cells expressing Na_V_1.3 labeled with 60 μM OptoDArG demonstrate the generation of depolarizing currents upon UV (60 mW) and hyperpolarizing currents upon blue light exposure (60 mW). The cell was held at −70 mV during the laser pulses; the duration of exposure is indicated by bars in the current trace. The current trace was recorded in the presence of 50 μM Gd^3+^ added to the external solution (Methods section). **d**, Method used for inferring changes in *C* upon photoisomerization: Under voltage-clamp, applying a voltage step, Δ*V*, to the membrane evokes a capacitive current. This current is given by Eq. 2 under the condition that d*C*/d*t*=0 and d*V*_s_/d*t*=0. Integration of this expression analogous to Eq. 8 results in *Q*_cap_=*C*×Δ*V*. Thus, *C* is inferred from a numerical integration of the capacitive current peak evoked upon a voltage step. **e**, Optocapacitive currents were generated by UV and blue light exposure; after these had decayed, voltage steps queried the instantaneous value of *C*, as described in **d**. Thus, the voltage step following UV light reveals *C* in the cis photostationary state, *C*_cis_, and following blue light *C* of the trans photostationary state, *C*_trans_. To obtain absolute values for *C*_cis_ and *C*_trans_, slow capacitance compensation was off. As in **c**, the displayed current trace was recorded in the presence of 50 μM Gd^3+^. **f**, The upper panel gives *C*_cis_ and *C*_trans_ determined as described under **e** for 10 different cells. As observed in PLBs, consistently *C*_cis_>*C*_trans_. The lower panel gives the corresponding percentages of change in *C*, (*C*_cis_−*C*_trans_)/C_cis_×100.

Subsequently, we sought to gain a rough estimate for the amount of membrane-embedded OptoDArG. We did so by capacitance measurements, measuring the time integral of the transient current for voltage steps (Fig. 3d). A measurement of *C* in the cis photostationary state, *C*_cis_, and the trans photostationary state, *C*_trans_, is shown in Fig. 3e. The result of measurements on 10 cells is shown in the upper panel of Fig. 3f. Subsequent calculation of the percentage change in *C* results in a distribution of values (lower panel in Fig. 3f), frequently around 5%. PLBs doped with 10 m% OptoDArG yielded similar changes in capacitance, suggesting that the cellular membranes may also have contained up to 10 m% OptoDArG.

### OptoDArG triggers photo-induced APs

As entailed by Eq. 2, optocapacitive currents modulate the membrane potential. To explore this, we did whole-cell recordings in current-clamp mode using HEK293 permanently transfected with Na_V_1.3. To initiate an AP is necessary only Na^+^ currents carried by Na_V_ channels ^39^, which will depolarize the cell membrane. In our case we exploit the endogenous HEK293 K_V_ and leak channels to repolarize the cell membrane. Therefore, the HEK293 Na_V_1.3 can be regarded as an artificial neuron regarding the generation of an AP. In HEK293 Na_V_1.3, exposure to UV light generated rapid optocapacitive depolarization (Fig. 4a), in agreement with the observation of depolarizing capacitive currents under voltage-clamp (Fig. 3c). Depolarization subsequently triggered the opening of Na_V_1.3 channels which generated an action potential (Fig. 4a). To the best of our knowledge, this is the first demonstration of direct photolipid-induced depolarization triggering an AP.

**Fig. 4:**
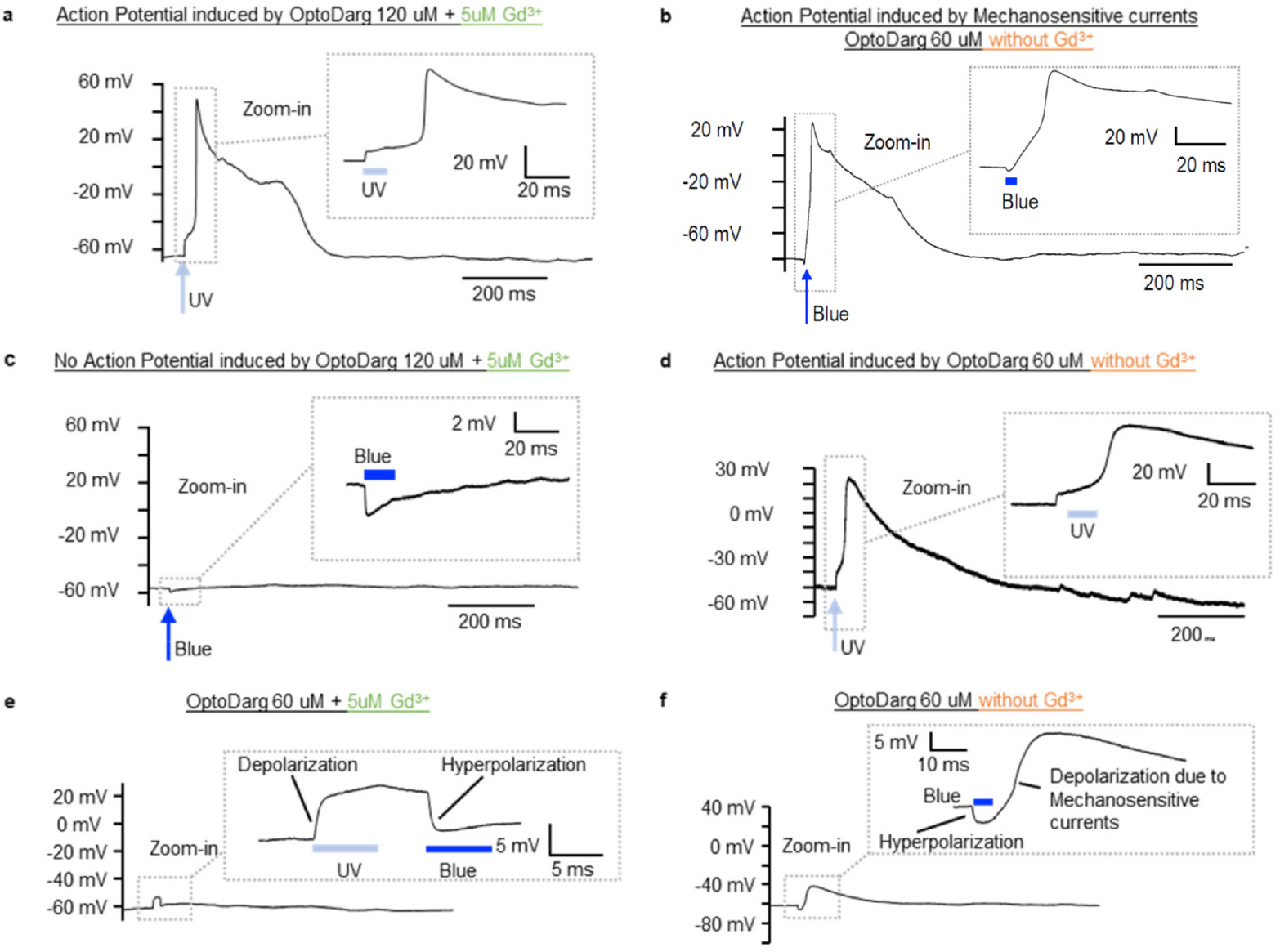
Photo-induced APs. The panels show current-clamp recordings of HEK293 Na_V_1.3 under varying conditions. UV laser power was 120–121 mW; blue laser power was 60 mW. **a**, AP elicited by OptoDArG-mediated optocapacitive depolarization upon UV light exposure recorded in the presence of 5 μM Gd^3+^; OptoDArG was 120 μM. The AP shown in **a** was recorded under conditions where the driving force for Na^+^ ions was augmented (see Methods). **b**, In the absence of Gd^3+^, hyperpolarization upon blue light exposure is followed by depolarization and AP generation; OptoDArG was 60 μM. Depolarization is attributed to mechanosensitive currents activated by the transition from cis-to trans-OptoDArG (see **f**). **c**, In the presence of 5 μM Gd^3+^ and 120 μM OptoDArG, blue light causes hyperpolarization, and neither depolarization nor AP generation. **d**, Same as **a**, but in a different cell, in the absence of Gd^3+^ and with 60 μM OptoDArG. **e**, OptoDArG-induced depolarization with UV followed by hyperpolarization with blue illumination. **f**, Blue light-driven hyperpolarization induced by cis to trans photoisomerization of OptoDArG and subsequent depolarization induced by mechanosensitive currents. Note that in some cases the depolarization generated by the photolipid currents did not reach the threshold to generate an AP, as shown in **e**. Similarly, in some cases, the depolarization induced by mechanosensitive currents did not reach the threshold to fire an AP, as shown in **f**.

Exposure to blue light evoked hyperpolarization, which was followed by depolarization and subsequent AP generation (Fig. 4b). Since the transition to trans-OptoDArG is associated with the generation of membrane tension (Fig. 2i), we added Gd^3+^, a broad inhibitor of mechanosensitive channels ^40, 41^. When repeating the experiment in the presence of 5 μM Gd^3+^, the large depolarizing currents were inhibited and photostimulation with blue light did no longer evoke APs (Fig. 4c). This differential response to blue light exposure between the absence (Fig. 4b) and presence of Gd^3+^ (Fig. 4c) was not due to Na_V_1.3 being blocked by 5 μM Gd^3+^ (Supplementary Fig. 4b); also, AP generation upon UV light exposure did not depend on the presence (Fig. 4a) or absence of 5 μM Gd^3+^ (Fig. 4d). We conclude that the rapid transition to trans-OptoDArG mediated by intense blue light triggered a Gd^3+^-sensitive depolarizing current, potentially facilitated by endogenous mechanosensitive channels ^42^.

Next, we studied the blue-light evoked capacitive and ionic currents under voltage-clamp conditions in cells and found further evidence for the contribution of mechanosensitive channels. Immediately following the hyperpolarizing optocapacitive current, another current emerges in the absence of Gd^3+^ (Fig. 5a). We find that this current is non-selective, reversing at around 0 mV (in sharp contrast to the optocapacitive current), its activation is not voltage-dependent, and it is sensitive to Gd^3+^. In the presence of 10 μM Gd^3+^ these ionic currents are subdued (Fig. 5b), as is blue light-triggered delayed depolarization (Fig. 4c). Most likely, they are facilitated by endogenously expressed mechanosensitive channels present in HEK293. We also observed the light-evoked mechanosensitive currents in blank HEK293T cells (not expressing Na_V_1.3), evidencing that the currents were not carried out by Na_V_1.3 (Supplementary Fig. 5) and suggesting that they are from an endogenous channel. Since HEK293 cells endogenously express a myriad of different ion channels ^43^, the mechanosensitive channel responsible for the hyperpolarizing currents observed here remains yet to be identified.

**Fig. 5:**
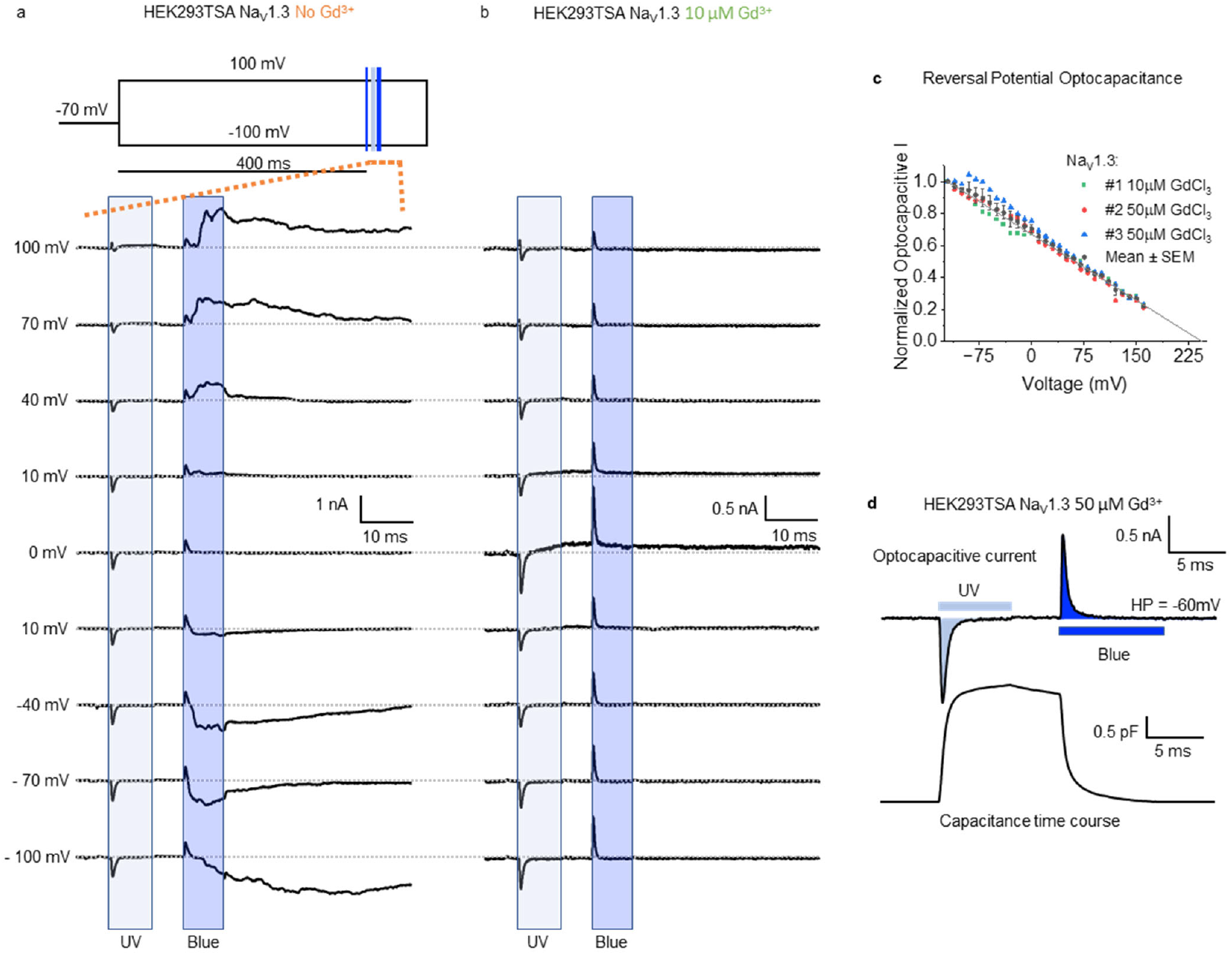
Voltage-clamp study on Gd^3+^-sensitive currents evoked upon switching to trans-OptoDArG. Panels **a** and **b** show voltage-clamp recordings of HEK293 Na_V_1.3. UV laser power was 97–100 mW; blue laser power was 60 mW. The voltage protocol for **a** and **b** is illustrated in the inset of **a**; the shown period of the obtained current traces is indicated by orange dashed lines. We used three laser pulses: (1) a blue pulse (3 ms) to achieve the trans photostationary state; (2) a UV pulse (6.5 ms) to photoisomerize trans-to cis-OptoDArG which generates a depolarizing current; (3) another blue laser pulse (6 ms) rapidly achieves trans-OptoDArG which results in a hyperpolarizing current. Both, depolarizing and hyperpolarizing capacitive currents in **a** and **b** reverse at large positive *V*, qualitatively consistent with thermally-evoked optocapacitive currents in cells. **a**, In the absence of Gd^3+^, the blue light-evoked optocapacitive current is followed by a current that reverses at around 0 mV. To avoid contamination of the mechanosensitive currents with the Na^+^ currents carried by Na_V_1.3, we applied a 400 ms depolarizing pulse to ensure that all sodium channels were in the inactivated state before evoking the mechanosensitive current by the laser pulses. **b**, In the presence of 10 μM Gd^3+^, the hyperpolarizing capacitive currents decay, and no additional currents are triggered. Cells in **a** and **b** are from different dishes. **c**, Graph of normalized blue light-evoked hyperpolarizing current amplitude over clamped voltage; a linear fit reveals the large positive reversal potential, *V*_rev_, of the optocapacitive current. **d**, Optocapacitive currents elicited by UV and blue light from a cell held at −60 mV and in the presence of 50 μM Gd^3+^. The time course of capacitance change was calculated according to Eq. 8 with *V*_rev_=*V*_s_.

Plotting the amplitude of the hyperpolarizing current vs holding voltage reveals a large positive reversal potential, *V*_rev_, in the range of +230 mV (Fig. 5c). It is recorded in the presence of Gd^3+^ as determining the reversal potential is otherwise hampered by the current generated by mechanosensitive channels. The potential is too large to be attributed to the intrinsic difference in HEK cell surface potential alone. Yet, Gd^3+^ may have contributed as its one- sided adsorption alters this difference profoundly ^8, 11^. Since the rate of change of capacitance (d*C*/d*t*) determines the effectiveness to evoke AP by the optocapacitive approach^10, 13^, the change in capacitance using photolipid occurred in less than (∼1.5ms - Fig. 5d) demonstrating the effectiveness of this method.

From the above experiments, we infer a molecular mechanism for depolarization following capacitance-decreasing and initially hyperpolarizing photolipid photoisomerization. Specifically, blue light gives rise to mechanical membrane tension. Mechanosensitive ion channels open in response to this stimulus, producing a depolarization that evokes an AP.

It is important to note that other factors may have contributed to the blue light-driven augmented cellular excitability in addition to tension. Since ion channels are susceptible to the physical state of the embedding matrix ^25^, the photo-induced changes in membrane material properties such as bilayer surface area and thickness ^17-19^, bending rigidity and area compressibility ^19, 44, 45^, as well as the propensity towards domain formation ^20, 22^ may also play a role. For example, changes in bilayer thickness and stiffness regulate the prominent bacterial mechanosensitive channels MscL and MscS ^23, 46^, and membrane mechanical calculations predict that bilayer bending rigidity contributes to Piezo1 gating ^47^.

Yet, whilst bilayer material properties are bound to affect mechanosensitive channels ^25^, the primary stimulus for their gating is stretch activation by membrane tension ^24^. Membrane tension – ‘force from lipids’ – opens the two-pore domain potassium ion channels TRAAK and TREK1 ^48^, the bacterial osmotic emergency valve MscL ^49^, as well as the eukaryotic stretch-activated channel Piezo1 ^50^. The effect of lipid transmitted tension must be distinguished from specific lipid mediated effects – as has been established for TREK1’s gating that may be enhanced by binding of anionic lipids ^51^.

## Conclusion

We propose a photolipid-based approach to induce direct depolarization and AP generation in excitable cells by light. The approach does not require the presence of intermediary mechanosensitive channels that would translate a light-induced increase in membrane tension into depolarizing currents. Further, it is easy to handle since externally added photolipids spontaneously insert into both leaflets of excitable cells, providing an elegant alternative method to genetic manipulations. Light power and photolipid concentration allow for adjustable modulation of the membrane potential.

## Methods

### Horizontal planar lipid bilayer experiments

*E. coli* Polar Lipid Extract (PLE) was obtained from Avanti Polar Lipids (distributed by Merck). OptoDArG was synthesized as described previously ^16^. Lipid aliquots and mixtures were prepared within amber glass micro reaction vessels from lipids dissolved in chloroform. Prior to storage at −80 °C, the solvent was evaporated by a mild vacuum gradient (Rotavapor, Büchi Labortechnik AG), and the dried lipids were flooded with argon.

Horizontal solvent-depleted planar lipid bilayers (PLBs) were folded from lipid monolayers on top of aqueous buffer in the lower and upper compartment of a custom-built chamber assembly made from PTFE ^20, 52^; the buffer used for all PLB experiments was 150 mM KCl, 10 mM HEPES, pH 7.4. First, an aperture between 70 to 100 μm in diameter in 25 μm-thick PTFE foil (Goodfellow GmbH) was created by high-voltage discharge. Following pretreatment of the septum with 0.6 vol% hexadecane in hexane, hexane was allowed to evaporate for >1 h, except for the experiment shown in Fig. 2d where the septum was applied shortly after pretreatment to enforce a large solvent torus. Then, the septum was attached by silicon paste to the lower side of the upper compartment of the chamber assembly. Lipid monolayers were prepared by applying lipid mixtures dissolved in hexane at a concentration of 10 mg/mL onto the aqueous interfaces of the upper and lower compartment. Unless noted otherwise, the lipid mixture was 80 m% *E. coli* PLE + 20 m% OptoDArG. After the hexane had evaporated, the folding of PLBs in a horizontal configuration was achieved by rotation of the upper compartment of the chamber assembly.

A 30 mm-diameter cover glass (No. 1, Assistent, Hecht Glaswarenfabrik GmbH & Co KG) fixed with a threaded PTFE ring comprised the bottom of the lower chamber. The chamber holder was installed on the sample stage of an Olympus IX83 inverted microscope equipped with an iXon 897 E EMCCD (Andor, Oxford Instruments Group). To position the horizontal PLB within working distance of a 40× magnification (1.30 NA) infinity-corrected plan fluorite oil immersion objective (UPLFLN40XO/1.30, Olympus), the chamber holder was equipped with screws for achieving fine translation of the upper compartment in z-direction. The real-time controller (U-RTC, Olympus) used for synchronizing lasers and electrophysiological acquisition, as well as the motorized microscope, were controlled using the proprietary cellSens software (Olympus).

For electrical measurements, Ag/AgCl agar salt bridges containing 0.5 M KCl were put into the compartments and connected to the headstage of an EPC 9 patch-clamp amplifier (HEKA Elektronik, Harvard Bioscience). The headstage and chamber assembly were housed in a Faraday cage. Voltage-clamp and software lock-in measurements were controlled using PATCHMASTER 2x91 software (HEKA Elektronik, Harvard Biosciences). In both recording modes, current was analogously filtered at 10 kHz by a combination of Bessel filters and acquired at 50 kHz. Sine wave parameters for software lock-in measurements (“Sine+DC” method with computed calibration) were 10–20 mV peak amplitude, 5 kHz, 10 points per cycle, no averaging; voltage offset was ±10 mV.

For cis to trans photoisomerization of OptoDArG-containing PLBs requiring blue light, the output beam of a 488 nm diode laser (iBEAM-SMART-488-S-HP, TOPTICA Photonics) was cleaned-up by a ZET488/10x excitation filter (Chroma), directed through a Keplerian-type beam expander with pinhole, and eventually focused into the back-focal plane of the objective via the IX83’s ZET488/640rpc main dichroic (Chroma). The diameter of the laser profile at the sample stage was ≈58 μm (1/e²). At a software-set output power of 200 mW, 30 mW exited the microscope objective, as determined by a photodiode (S120VC, Thorlabs).

For trans-to-cis photoisomerization requiring UV light, the output beam of a 375 nm diode laser (iBEAM-SMART-375-S, TOPTICA Photonics) was expanded using a separate Keplerian-type beam expander, focused into the back-focal plane of the objective via a ZT458RDC (Chroma) and the IX83’s ZET488/640rpc main dichroic. The diameter of the laser profile at the sample stage was roughly 150–250 μm; ≈30 mW exited the microscope objective at a software-set output power of 70 mW.

Electrophysiological data recorded with PATCHMASTER were exported and analyzed using Mathematica (Wolfram Research); the presented graphs were created in OriginPro 2023b (OriginLab Corporation). Integration of optocapacitive currents obtained at different clamped voltages and subsequent linear fitting was done by default to infer capacitance change and voltage offset (e.g., Figs. 2g and 2h); recordings indicating a voltage offset >±20 mV for symmetric bilayers were excluded from further analysis.

### Patch-clamp experiments

Whole-cell patch clamp experiments were performed according to Hamill, Marty, Neher, Sakmann and Sigworth ^53^. We used an in-house designed and built patch clamp amplifier for both current and voltage-clamp experiments. Data was sampled at 1 MHz, digitally filtered at Nyquist frequency, and decimated for the desired acquisition rate. To acquire the data, we used in-house software (Gpatch64MC) to control the 16-bit A/D converter (USB-1604, Measurement Computing, Norton, MA). The current and voltage signals were sampled at 100 kHz and filtered at 5 kHz. The current signal was filtered using a 4-pole Bessel filter (FL4, Dagan Corporation, Minneapolis, MN) and the voltage signal was also filtered using a 4-pole Bessel filter (Model 900, Frequency Devices, Haverhill, MA). Borosilicate patch pipettes (1BF120F-4, World Precision Instruments, Sarasota, FL) were pulled on P-2000 (Sutter Instruments, Novato, CA) horizontal puller, and the resistance ranged from 2-5 MΩ when back-filled with pipette solution. The pipette and bath solutions were based on Carvalho-de-Souza, Treger, Dang, Kent, Pepperberg and Bezanilla ^9^ and their composition were (mM): The internal (pipette) solution NaCl 10, KF 130, MgCl_2_ 4.5, HEPES 10, EGTA 9, and pH 7.3 (KOH) and the external solution NaCl 132, KCl 4, MgCl_2_ 1.2, CaCl_2_ 1.8, HEPES 10, glucose 5.5 and pH 7.4 (NaOH). In some cases, to increase the driving force for Na^+^ ions, we replaced NaCl from the pipette solution to KCl, and increased NaCl on the external to 140 mM. Recordings were performed at room temperature (≈17–18 ºC). When necessary, we used solutions containing GdCl_3_ (439770, Sigma-Aldrich, St. Louis, MO) from 30 mM stock solutions dissolved in water. This stock solution is then diluted to the bath solution at the desired final concentration. Analysis was performed using in-house software. Analysis and graphs were constructed using OriginPro 2023b (Origin Lab Corporation, Northampton, MA, USA).

### Optical Stimulation setup

We used a Ti : Sapphire (Mai Tai HP, Spectra-Physics, Milpitas, CA) laser combined with a multi-harmonic generator (ATsG-3-0.8-P, Del Mar Photonics, San Diego, CA) to obtain 367 nm laser wavelength for the UV light and the blue light we used a laser diode 447 nm (ams OSRAM) mounted on a cage (CP1LM9, Thorlabs, Newton, NJ) with aspheric lens (LDH9-P2, Thorlabs). The maximum power for the UV light at the objective output was around 100 mW, and for the blue laser was 1 W. We used a camera (DCC1545M-GL, Thorlabs) mounted at an in-house built microscope to visualize the cells. To allow maximum UV light transmission, we used a fused silica lens designed in-house. We used a shutter (VS25S1ZO, Vincent Associates, Rochester, NY) to get onset laser pulses in less than 15–20 μs. We used custom-designed electronic circuitry to achieve fast onset of laser pulses with the blue laser diode.

### Cell labeling

To label the HEK293 cells with the OptoDArG we followed the protocol established by Leinders-Zufall, Storch, Mederos, Ojha, Koike, Gudermann and Zufall ^54^. Briefly, prior to labeling, the stock solution containing 60 mM of OptoDArG, dissolved in dimethyl sulfoxide (DMSO - 276855, Sigma-Aldrich), was heated for 5 minutes at 37 ºC. The stock solution was diluted with recording/bath solution to the final concentration of 60 μM. In some cases, to increase the amount and efficiency of labeling, we used 120 μM OptoDArG (indicated in the text when used). The solution was then vortexed and incubated at 37 ºC for 5 minutes. The dish containing the cells was taken from the incubator and washed 1–3 times with the external solution. After washing the dish containing the cells, we added the solution containing OptoDArG and incubated it at 37 ºC for 1 hour. After labeling, we washed the dish 1–3 times with the external solution and assembled it in the recording chamber. The stock solution containing the OptoDArG was aliquoted in small volumes (≈6 μl) and stored at −20 ºC. The DMSO concentration never exceeds 0.2% in the experiments.

### Cell Culture

HEK293 cells stably transfected with Na_V_1.3 channels were purchased from American Type Culture Collection (CRL-3269, ATCC, Manassas, VA). They were cultured on 75 cm^2^ flasks in a humidified incubator at 37 ºC with 5% CO_2_, following ATCC protocols. The cells were grown and maintained in DMEM/F12 medium (ATCC 30-2006) supplemented with 10% heat-inactivated fetal bovine serum (FBS - SH30071.03, HyClone-GE healthcare), 0.5 mg/ml G-418-sulfate (10131-027, Gibco, ThermoFisher), 100 U/ml penicillin-G sodium and 100 μg/ml streptomycin sulfate (P4333, Sigma-Aldrich) and 2.0 μg/mL Puromycin (A11138-03, Gibco). Dr. Eduardo Perozo kindly provided HEK293T cells, and they were grown in Dulbecco’s modified Eagle’s medium (DMEM - 11995065, Gibco) supplemented with 10% FBS and 100 U/ml penicillin-G sodium and 100 μg/ml streptomycin sulfate. They were cultivated at 37 ºC in 5% CO_2_ humidified incubator. Before experiments, cells were detached using Accutase (AT-104, Innovative Cell Technologies, Inc., San Diego, CA) following the manufacturer’s protocol. They were centrifuged for 5 minutes at 125 ×g; the supernatant was replaced by fresh medium and seeded (≈30-50% confluency) into previously prepared Poly-L-lysine treated culture dishes. They were used within 12–36 h for electrophysiological experiments. Glass-bottomed culture dishes (D35-10-1-N, Cellvis, Mountain View, CA) were incubated with Poly-L-lysine solution (P8920, Sigma-Aldrich) for 15 min at room temperature and thoroughly rinsed with sterile PBS (10010-023, Gibco) and stored until use.

## Acknowledgments

We want to thank Drs. Navid Bavi and Eduardo Perozo for their insightful discussions and suggestions. This work is supported by the Austrian Science Fund (FWF) Award P34826 to PP, National Institutes of Health Award R01GM030376 to FB, and National Science Foundation Award QuBBE QLCI (NSF OMA-2121044) to FB.

## Author Approvals

All authors have seen and approved the manuscript. It has not been accepted or published elsewhere.

## Competing Interests

The authors declare no competing interests.

## Notes

### Competing Interest Statement

The authors have declared no competing interest.

