## Supplement for "Photolipid excitation triggers depolarizing optocapacitive currents and action potentials"

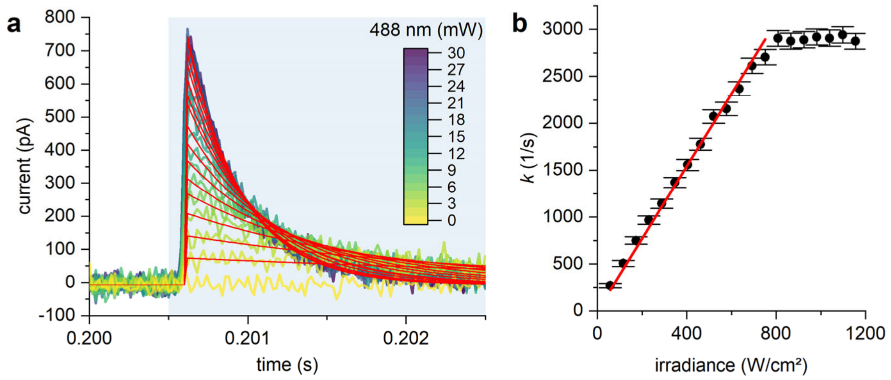


**Supplementary Fig. 1: Power-dependence of $I_{cap}$ decay rate. a,** The power-dependent $I_{cap}$ upon blue light exposure shown and described in Fig. 1e were fitted with a monoexponential decay model of the form: $y(t)=y_0$ for $t\leq0.2006$ s and $y(t)=A\times\exp(-(t-0.2006)\times k)+y_0$ for $t>0.2006$ s. A slight delay of ≈100 µs was introduced between the onset of light exposure and the start of exponential fitting (0.2006 s) to avoid the initial rise of the current which is not fully resolved due to analog filtering. **b,** The apparent rates of decay from the exponential fits in **a**, *k*, are plotted over irradiance estimated from power at the sample stage divided by area calculated from 1/e$^2$-diameter of the blue laser profile. The data is fit between 60 and 750 Wcm$^{-2}$ by a linear model of the form $y(x)=a\times x$, whereby $a$=3.85 cm$^2$W$^{-1}$s$^{-1}$ ($R^2>0.99$).

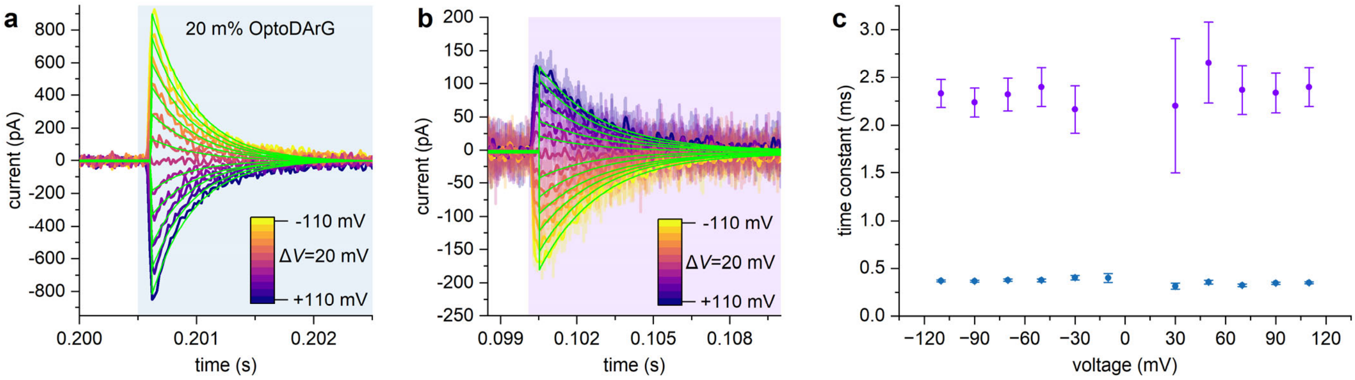


**Supplementary Fig. 2: The rate constants of $I_{cap}$ decay are voltage-independent, in accordance with the theoretical prediction. a,b,** $I_{cap}$ shown in Figs. 1c and 1d were fit as described in Supplementary Fig. 1a. For UV light-evoked $I_{cap}$, the exponential decay was fit starting from 0.1005 s. **c,** The resulting time constants, $1/k$, are plotted over applied voltage. Purple points correspond to UV light-evoked $I_{cap}$ (**a**), blue points to blue light-evoked $I_{cap}$ (**b**). Differences in the decay rates are due to differences in irradiance (UV light irradiance was lower than blue light irradiance) and differences in photoisomerization rate for the cis to trans and trans to cis transition (Arya, Jelken et al. 2020). As expected from Eq. 7, the rate of decay does not depend appreciably on the applied voltage.

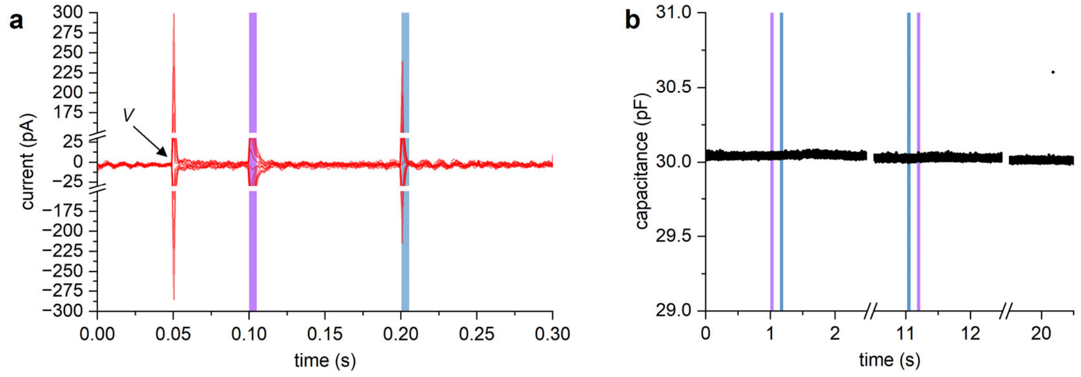


**Supplementary Fig. 3: Controls. a,** OptoDArG photoisomerization does not alter PLB conductance. The red traces are digitally filtered versions of the raw current traces from which Figs. 1c and 1d were prepared (the color coding according to power was omitted). As in Fig. 1, purple and blue background designate UV and blue light exposure, respectively. The arrow annotated *V* designates the application of voltage ranging from −110 mV to +110 mV with $\Delta V$=20 mV, as described in the caption to Fig. 1; prior to 50 ms, *V* was 0 mV. Note that the voltage jump- and photoisomerization-evoked capacitive currents are truncated to allow for a more focused view of the baseline. **b,** In the absence of OptoDArG, capacitance does not change upon blue or UV light exposure. The recording was done as in Fig. 2a but on a PLB folded from 100% *E. coli* PLE.

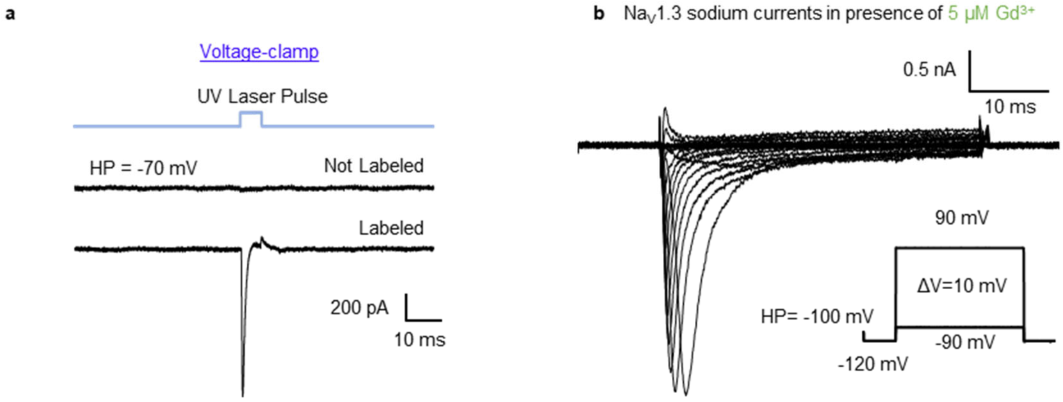


**Supplementary Fig. 4: No optocapacitive current is present when cells were not labeled with OptoDArG and 5 µM Gd$^{3+}$ does not affect Na$^{+}$ currents through Na$_V$1.3. a,** Without OptoDArG labeling, no optocapacitive currents were induced by UV light exposure, indicating the absence of photothermal effects due to UV light itself. **b,** Na$^{+}$ currents elicited by the voltage protocol inset in **b,** and in presence of 5 µM Gd$^{3+}$.

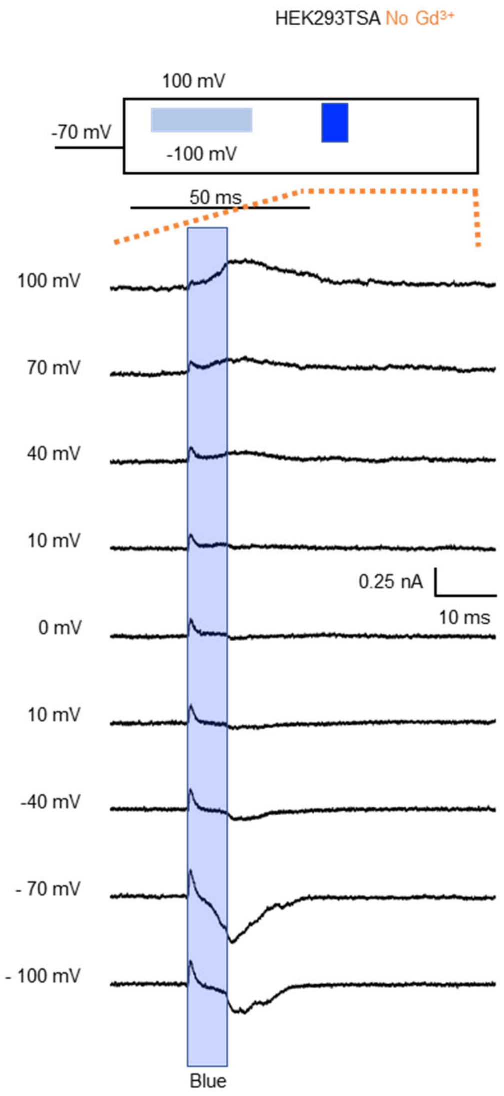


**Supplementary Fig. 5: Mechanosensitive currents evoked upon switching to trans-OptoDArG in blank HEK293T cells.** Currents were elicited using non-transfected (blank) HEK cells at different voltage. A 30 ms UV laser pulse, indicated as a light blue square (30 mW), was applied before to prime the cis to trans transition induced by the blue laser (160 mW), indicated as blue square in the voltage protocol inset. The voltage protocol is illustrated in the inset and the time shown is indicated by orange dashed lines.
